# Stoichiometric modeling of artificial string chemistries

**DOI:** 10.1101/2020.09.16.300491

**Authors:** Devlin Moyer, Alan R. Pacheco, David B. Bernstein, Daniel Segrè

**Affiliations:** Bioinformatics Program, Boston University, Boston, MA 02215; Department of Biology, Boston University, Boston, MA 02215; Biological Design Center, Boston University, Boston, MA 02215; Department of Biomedical Engineering, Boston University, Boston, MA 02215; Department of Physics, Boston University, Boston, MA 02215

## Abstract

Uncovering the general principles that govern the architecture of metabolic networks is key to understanding the emergence and evolution of living systems. Artificial chemistries, *in silico* representations of chemical reaction networks arising from a defined set of mathematical rules, can help address this challenge by enabling the exploration of alternative chemical universes and the possible metabolic networks that could emerge within them. Here we focus on artificial chemistries in which strings of characters represent simplified molecules, and string concatenation and splitting represent possible chemical reactions. We study string chemistries using tools borrowed from the field of stoichiometric constraint-based modeling of organismal metabolic networks, through a novel Python package, ARtificial CHemistry NEtwork Toolbox (ARCHNET). In addition to exploring the complexity and connectivity properties of different string chemistries, we developed a network-pruning algorithm that can generate minimal metabolic networks capable of producing a specified set of biomass precursors from a given assortment of environmental molecules within the string chemistry framework. We found that the identities of the metabolites in the biomass reaction wield much more influence over the structure of the minimal metabolic networks than the identities of the nutrient metabolites — a notion that could help us better understand the rise and evolution of biochemical organization. Our work provides a bridge between artificial chemistries and stoichiometric modeling, which can help address a broad range of open questions, from the spontaneous emergence of an organized metabolism to the structure of microbial communities.

## Introduction

Metabolism occupies a central role in the functioning of biological systems, yet much remains unclear about the degree to which basic features of metabolic networks reflect evolutionary accidents or optimal network structures [1–4]. In parallel to analyses focused on metabolism as we know it in individual organisms [5,6] or in the whole biosphere [2,7,8], multiple studies have explored the utility of abstract models of chemistry to investigate particular features of chemical networks. These models, also known as artificial chemistries, have the benefit of being unconstrained by the limits of what is known about extant metabolism and about its possible intermediate states lost through evolutionary history [9–11]. Artificial chemistry has been used to study various aspects of the origin of life from abiotic chemistry [9,11,12], common structural features of metabolic networks (e.g. hub metabolites) [13–15], the general behavior of chemical (not necessarily biochemical) reaction networks [10,16], the optimality (or lack thereof) of metabolic networks [17,18], among other questions [9]. The artificial chemistry models used in these studies typically employ highly abstracted representations of chemistry [9,11,17]. However, more precise and realistic models involving either string rules based on formalization of real chemistry (like SMILES [19] and variants thereof [20,21]), or *de novo* approximate quantum mechanics computations [10], have been used to explore the full space of possible real-life chemistry up to a certain degree of complexity [22]. Artificial chemistry approaches have yielded many insights into general features of metabolism, but these findings have remained largely disconnected from the large body of metabolism research focused on characterizing real metabolic networks. We believe that many novel insights into metabolism will be enabled by combining artificial chemistry with techniques commonly used to study real metabolic networks.

The field of stoichiometric constraint-based modelling has provided many approaches that can be particularly useful for quantitatively understanding the structure and function of metabolic networks [23–26]. In particular, Flux Balance Analysis (FBA) is a common technique for studying metabolic networks at the level of a whole organism. FBA estimates the space of possible fluxes through the network at steady-state, and is generally employed to identify metabolic regimes closed to a biologically meaningful optimum [27]. FBA has been used to simulate multiple types of experiments and phenotypes, such as growth rates and metabolic phenotypes of gene knockouts, growth efficiency on different media, and identification of potential drug targets [27–29]. While FBA and stoichiometric constraint-based modeling have been widely used on real organism’s metabolic networks, these techniques have only rarely been applied to artificial chemistry networks.

In the present work, we use a specific type of artificial chemistry known as a string chemistry, where each molecule is represented by a string of characters (Figure 1) [9,11,17]. Our string chemistry model is relatively simple: all strings (i.e. metabolites) are linear sequences of characters (i.e. monomers, atoms, functional groups) that may react by either concatenating end-to-end or splitting into two smaller strings (see Methods). A particular string chemistry network is defined by the set of different characters each metabolite can be composed of and the maximum length a metabolite can reach. While these rules are much simpler than those governing real chemical reactions, Riehl et al. managed to find structural similarities between real metabolic networks and string chemistry networks with only one type of character (i.e. the only difference between any two metabolites is their length) [17], so we expect that string chemistries chemistry with more than one monomer type may yield further insights into the general properties of metabolic networks.

**Figure 1.**
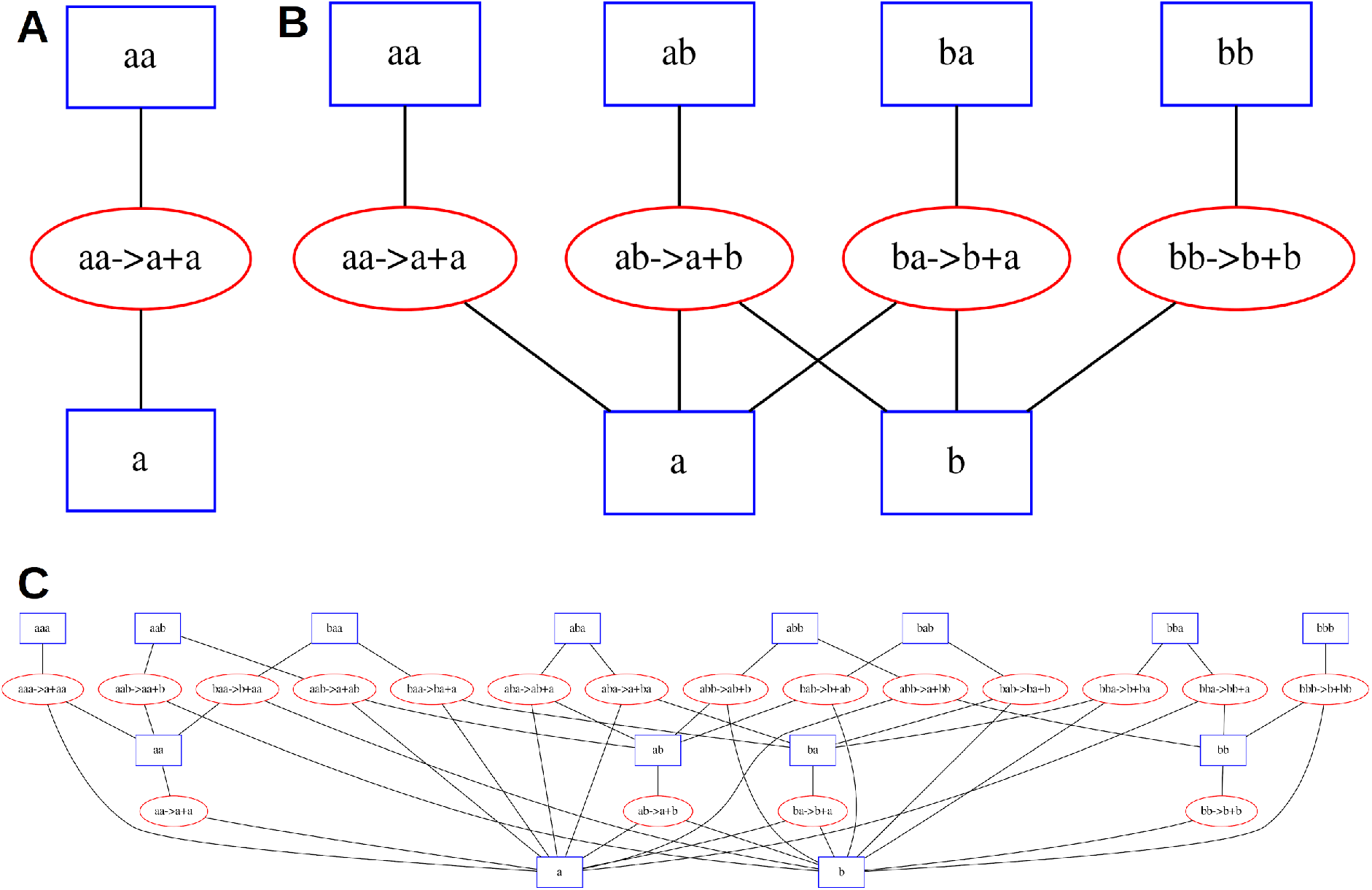
Three simple string chemistry networks. Square nodes represent chemicals and oval nodes represent reactions. Edges connect chemicals to the reactions they participate in, either as reactants or products. **A**. A network with only one type of monomer and a maximum string length of 2. **B**. A network with two types of monomers and a maximum string length of 2. **C**. A network with two types of monomers and a maximum string length of 3.

In this manuscript, we describe the ARtificial CHemistry NEtwork Toolbox (ARCHNET), a Python package we created for generating string chemistry networks of arbitrary size and implementing stoichiometric modeling algorithms (including FBA) on those networks. Using this string chemistry framework, we created an algorithm for determining the minimal metabolic network capable of producing a given set of metabolites (“biomass precursors”) from another set of metabolites (“environmental nutrients”). Our analysis of random choices of nutrients and biomass precursors in different string chemistry networks provides new insight into the rules governing the structures of these minimal metabolic networks and suggests possible implications for the study of real metabolic networks.

## Methods

### Artificial Chemistry Model

The artificial chemistry model used here is an extension of the one used in [17] and is similar to previously used artificial chemistries (e.g. [9,11,14]): each “chemical” is a string of characters of some arbitrary length, where each character represents an individual atom (or functional group, or monomer). A chemical may condense with one other chemical to produce a longer chemical; the two strings are simply concatenated (e.g. ab + aa → abaa). A chemical may also split into two smaller chemicals at any point along its length (e.g. ababb → ab + abb). Only pairwise condensation/dissociation reactions were considered due to the rarity of termolecular and higher reactions in real chemistry [30–32]. For simplicity, all reactions are modeled as being completely reversible, even though in principle further constraints on reversibility could easily be added. The numbers of chemicals and allowed reactions in the model are functions of the number of unique characters (“monomers”) and the maximum chemical length. These functions can be obtained analytically by enumerating the sizes of various string chemistry networks and examining the resulting series:

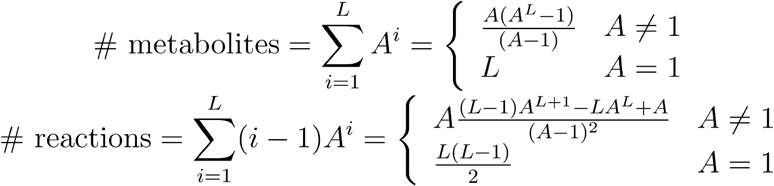

where *A* is the number of unique characters (monomers) and *L* is the maximum chemical length. We will refer below to a specific complete set of metabolites and reactions generated for a given choice of *A* and *L* as a “chemical universe”. This will allow us to clearly distinguish such complete sets from subsets generated by pruning algorithms (see below).

### Flux Balance Analysis

Flux Balance Analysis (FBA) is a mathematical framework for computing steady-state fluxes through chemical reactions in a given network of reactions subject to linear constraints [27]. The network of reactions is represented as a stoichiometric matrix **S**, where each column contains all of the stoichiometric coefficients for an individual reaction (negative for substrates, positive for products) and each row indicates how much of an individual metabolite is produced or consumed by each reaction (see Figure 3 for an example of a string chemistry network and its associated stoichiometric matrix). The reaction fluxes to be computed are represented by a vector **v**. In order for the network to be at steady-state, **v** must be in the null space of **S**:

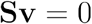

The resulting system of equations is underdetermined for nearly all nontrivial networks. Additional constraints may be specified that limit the values of fluxes through specific reactions, typically reflecting known thermodynamic constraints on certain reactions. These constraints typically reduce the space of feasible solutions, but still leave the problem underdetermined. Thus, a linear combination of reactions **Z** (the objective function) through which flux should be maximized is also typically specified:

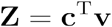

where **c** is a vector indicating which reactions are to be included in the objective function **Z**. As FBA is usually applied to biochemical reaction networks, the objective function frequently is set to correspond to a single reaction that produces the right proportion of all precursors necessary for the generation of cellular macromolecules and key metabolites, representing growth of cellular biomass. While FBA was originally developed for studying and engineering microbial metabolic networks, its formalism is easily adaptable to any chemistry, provided that its chemical reactions can be represented as columns of a stoichiometric matrix (Figure 3).

### The ARtificial CHemistry NEtwork Toolbox (ARCHNET) Package

We created a Python package to facilitate the creation and handling of string chemistry networks (as defined above) of arbitrary size and the application of FBA to such networks. All FBA computations were performed using the COBRApy Python package [24]. This package, along with all scripts used to generate data and create figures are available in a GitHub repository: https://github.com/segrelab/string-chemistry. The package contains tools for generating and analyzing string chemistry networks of arbitrary size, given the set of characters to use as monomers and the maximum string length. The networks can be returned as a stoichiometric matrix and/or a COBRApy model (to facilitate doing FBA or any other stoichiometric modelling technique).

### Network Pruning Algorithm

We implemented an algorithm that takes a complete string chemistry network as an input (e.g. the network of all possible reactions and metabolites when there are *A* = 2 and *L* = 5), and generates as an output a subnetwork that has been pruned to satisfy specific criteria. Specifically, given (i) a string chemistry network, (ii) a biomass composition (i.e. a set of molecules that have to be produced at stoichiometrically fixed proportions) and a (iii) set of available environmental resources, the algorithm iteratively removes reactions from the network until there is no flux through the output reaction (Figure S1). In particular, it repeatedly runs FBA to assign fluxes to all reactions and removes reactions with no flux and the reaction with the smallest nonzero flux. Once there is no flux through the output reaction, the last reaction that was removed is added back to the network and the network is “pruned”. The pruning algorithms are part of the Python package described above. Several other assorted scripts provide examples of applications of this pruning algorithm to string chemistry networks.

## Results

### A Python Package for Creating and Analyzing Arbitrary String Chemistries

We have created the ARtificial CHemistry NEtwork Toolbox (ARCHNET), a Python package capable of generating string chemistry networks of arbitrary sizes given the number of unique characters (*A*) and the maximum length of a string (*L)* (Figure 1). For simplicity, the only types of reactions allowed in these networks are pairwise string concatenation and splitting (see Methods for more details). Even with this restriction on reaction complexity, the networks increase in size very rapidly as *A* and/or *L* increase (Figure 2AB and Methods). For example, a basic chemistry with *A* = 3 and *L* = 2 would have 12 metabolites and 9 reactions. If we increase *A* by 1, the network would involve 20 metabolites and 16 reactions. If we instead increased *L* by one, there would be 39 metabolites and 63 reactions. Clearly, the network sizes depend very differently on these two parameters (see Methods). One of the important features of the package is that it can output networks both as a simple text file containing the stoichiometric matrix and as a COBRApy model [24], which can be exported as an SMBL file [33], so most tools developed to study real metabolism, including standard FBA calculations (Figure 3), are straightforward to apply to our string chemistries.

**Figure 2.**
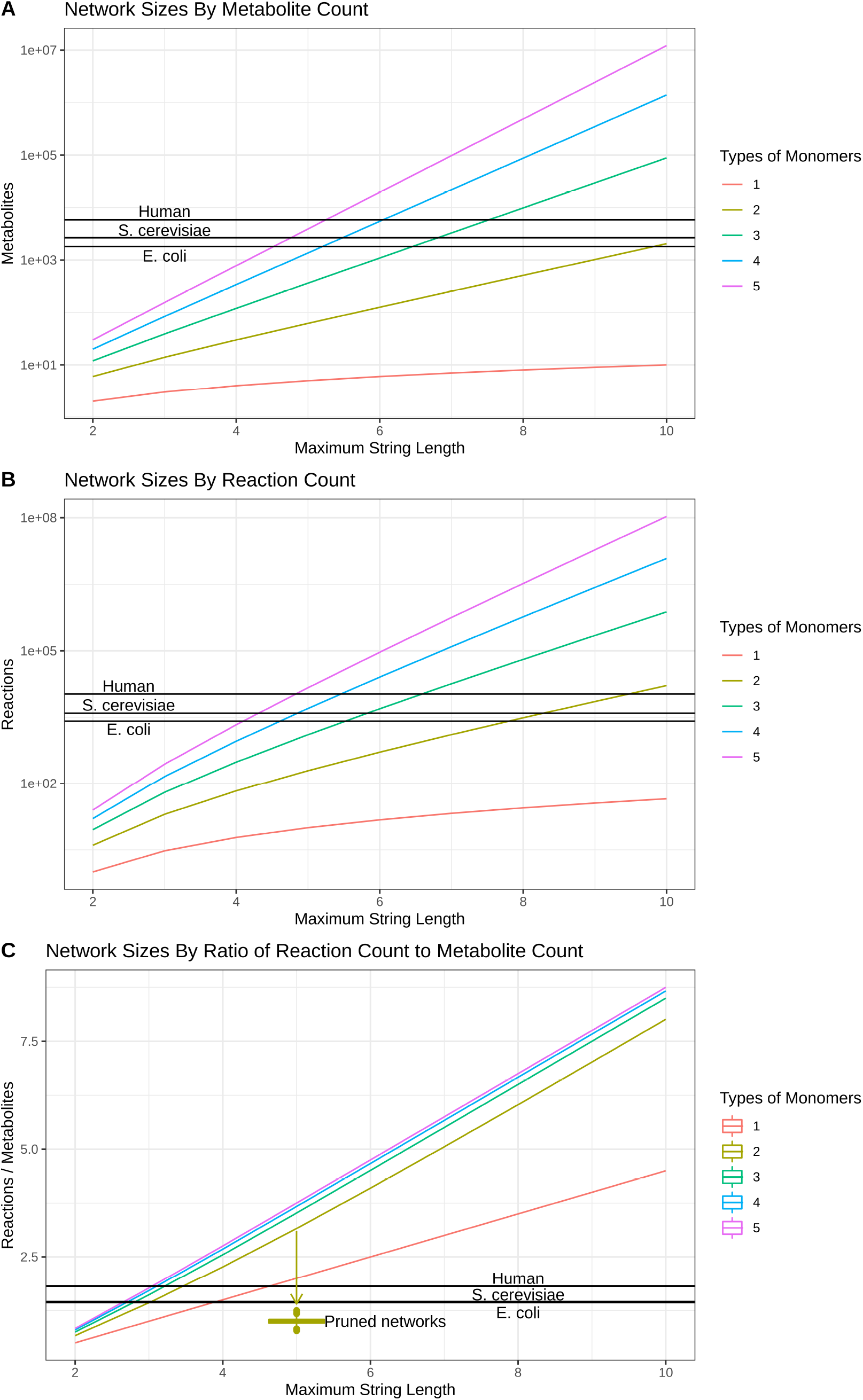
Comparison of size and connectivity of string chemistry networks (colored lines) to real metabolic networks (black lines). **A**. Network sizes measured by metabolite counts. **B**. Network sizes measured by reaction counts. **C**. Network connectivities measured by ratio of reactions to metabolites. Ratios for networks pruned from the chemical universe with *A* = 2 and *L* = 5 are shown as a boxplot.

**Figure 3.**
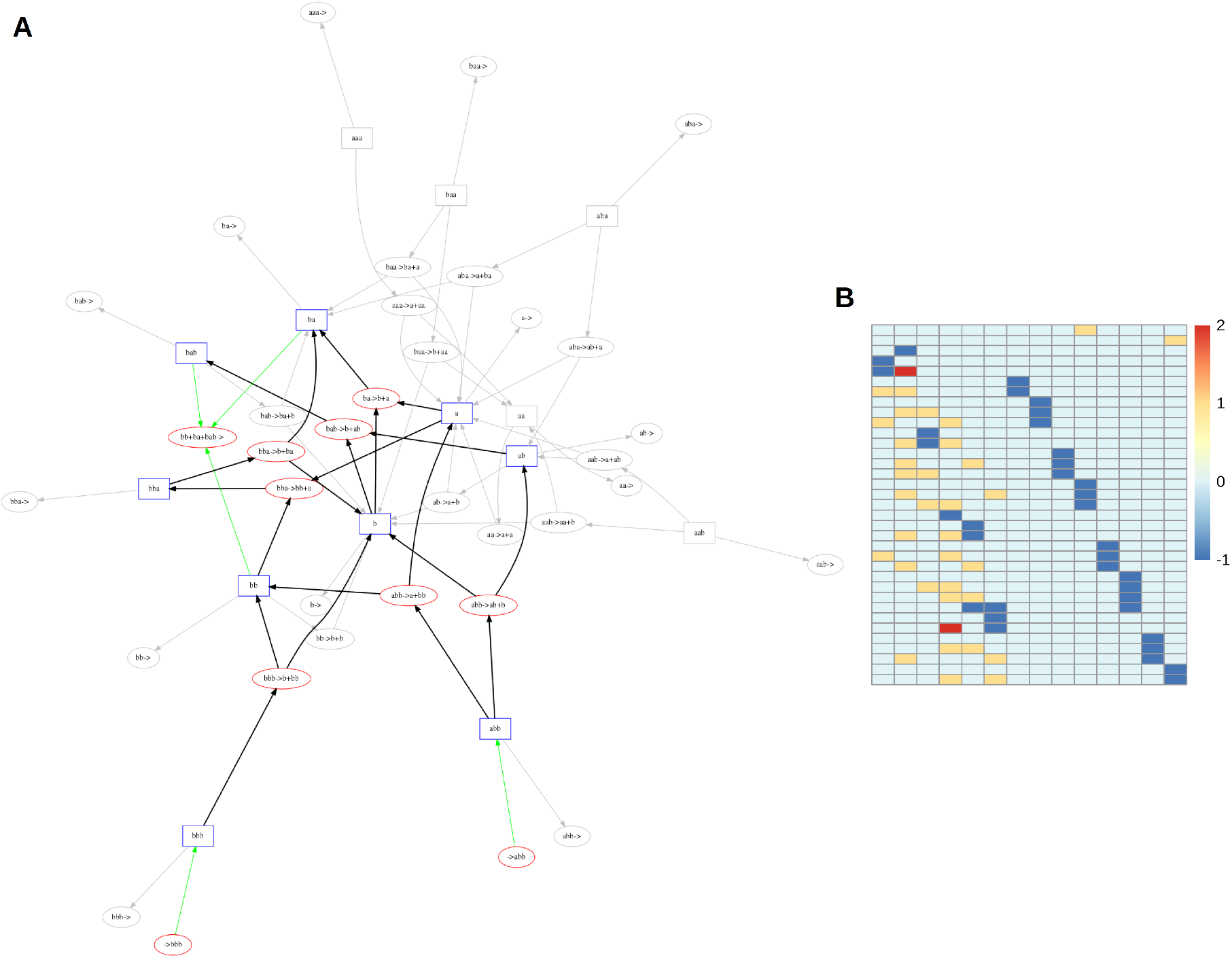
Flux-Balance Analysis on string chemistry networks. **A**. String chemistry network with A = 2 and L = 3. Metabolites are represented by blue rectangles and reactions are represented by red ovals. Edge colors represent reaction fluxes after maximizing flux through the biomass reaction: green edges are exchange fluxes (import/export/biomass production), black edges represent nonzero fluxes, and grey edges represent fluxes of zero. The direction of non-grey edges corresponds to the direction of flux; directions on grey edges are arbitrary. **B**. Stoichiometric matrix of network in **A**.

While real metabolic networks are much more complex than our string chemistry networks, both in terms of the underlying molecular structures and of the possible types of reactions, there are specific metrics that can be computed both for real metabolic networks and for our string chemistry networks. By comparing such metrics between these two systems, one can appreciate that artificial chemistries, despite their apparent simplicity, can quickly approach sizes and complexities comparable to those of real metabolic networks (see also [17]). By evaluating (numerically or analytically; see Methods) the numbers of metabolites and reactions in various string chemistry networks, we see that even string chemistry networks with few unique characters and short maximum lengths (e.g. *A* = 4, *L* = 5; *A* = 2, *L* = 10) reach sizes comparable to those of the human, yeast and *E. coli* metabolic networks (Figure 2A,B). However, as seen in Figure 2C, these artificial networks have a much higher connectivity (ratio of reactions to metabolites) than the real organisms’ metabolic networks. Conversely, a simple network with *A* = 1 and *L* = 3 would already reach a connectivity similar to the one of real metabolic networks, but would obviously be much smaller. One could then ask whether it is possible to create string chemistry networks that are both of similar size and connectivity to those of real metabolic networks. Indeed, one should view the complete string chemistries depicted here as analogous to “complete chemical universes”, out of which a single organism’s metabolic network would constitute a small subset. As shown below, this concept can be explored in artificial chemistries by devising algorithms that can prune complete chemical networks to obtain subnetworks that resemble individual organisms’ metabolic networks.

### Pruned Networks as Proxies for Evolved Organisms

After demonstrating the capacity of our package to create stoichiometric matrices usable by standard modeling tools and comparing properties of string chemistry networks to real metabolic networks, we explored the properties of string chemistry subnetworks that more closely resemble the metabolic networks of individual organisms. We modeled organism-scale metabolic networks as “minimal” (i.e. using the fewest reactions) networks capable of producing a given set of metabolites (i.e. biomass precursors), which is consistent with a simple parsimonious evolutionary assumption. To identify these minimal networks, we implemented a “pruning” algorithm that iteratively applies FBA to string chemistry networks. Briefly, the algorithm works by running FBA on a string chemistry network (initially set to the whole chemical universe given particular values of *A* and *L*) with some specified nutrient uptake reactions (i.e. generate individual metabolites from nothing) and a “biomass” reaction (i.e. consuming specific —in this case, equal— ratios of a given set of metabolites), removing all reactions that have no flux, testing whether or not the reaction with the smallest nonzero flux can be removed without eliminating flux through the biomass reaction, and repeating until no reactions can be removed (see Methods and Figure S1). We explored two variants of this pruning algorithm: one that allowed “export” of any metabolic reaction as a waste product throughout the pruning process (i.e. each metabolite has a reaction that consumes it and produces nothing and it is never removed during the pruning process) and one that did not allow any metabolites other than the biomass precursors to be “exported”. Pruned networks tend to have slightly fewer reactions when all metabolites can be exported than when there are no export reactions, due to the fact that any excess metabolite can be secreted (similar to costless byproducts predicted to be secreted in real metabolic networks [34]), rather than recycled internally.

### Biomass Precursors Shape Network Architecture More Than Environmental Composition

Using this pruning algorithm, we investigated the relative importance of the choice of nutrients and the choice of biomass precursors on the structure of pruned networks. We generated the string chemistry universe with *A* = 2 and *L* = 5, then created different biomass compositions (100 different sets of 5 randomly-chosen biomass precursors) and different sets of nutrients (100 random pairs of nutrients) using the metabolites contained within this chemical universe. Note that upon choosing the biomass composition, a growth flux was added to produce all chosen biomass precursors in equal proportions (see Methods). We then ran the pruning algorithm on all possible combinations of these nutrients and biomass precursors (Figure 4A). In order to compare the structures of the pruned networks, each network was represented as a binary vector with as many elements as there were reactions in the chemical universe. In this binary vector, a 1 represents a reaction that was kept in the pruned network and a 0 represents a reaction that was removed during pruning. These binary vectors were visualized on UMAP [35] plots (Figure 4B-G). The main outcome of this analysis is that, regardless of whether or not export reactions are allowed, networks with the same biomass reaction typically cluster together, while networks with the same nutrient sources frequently have very different structures (Figure 4B, C). The clustering is generally a bit weaker in the networks pruned without export reactions—there are more isolated networks and distinct small clusters—but the pruned networks still noticeably by biomass reaction (Figure 4E, F). Note that the clustering of networks doesn’t seem to display any clear pattern in terms of achievable growth rates (Figure 4D,G), which are highly variable and roughly distributed around an intermediate value between zero and the maximum. In other words, networks with similar architecture, as dictated by the biomass composition, may achieve substantially different growth rates, suggesting that while biomass composition dictates network structure, environmental constraints affect efficiency of production.

**Figure 4.**
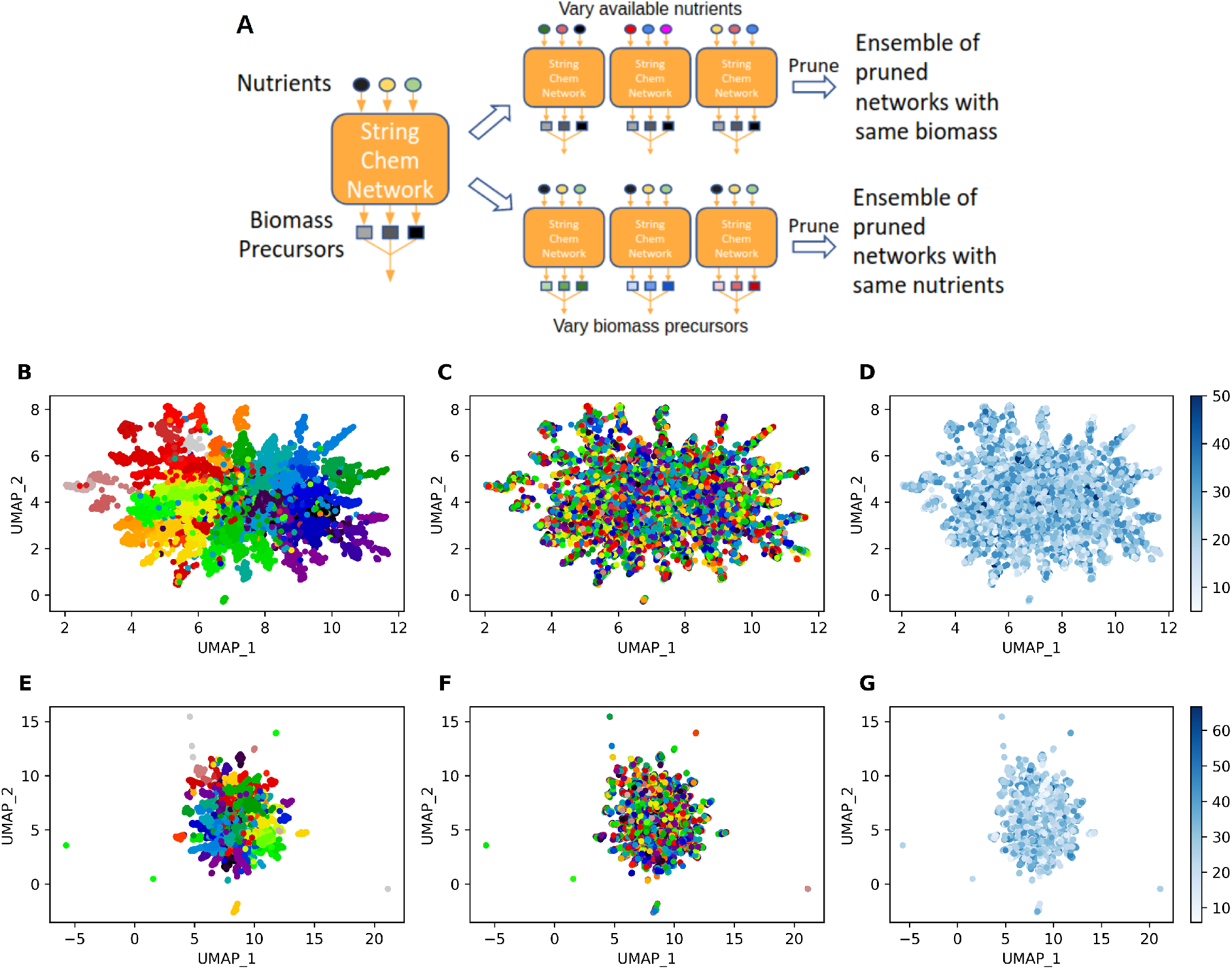
Choice of biomass precursors impacts structure of pruned networks more than choice of available nutrients. **A**. Cartoon representation of how data shown in panels **B**-**G** were generated. **B**. UMAP scatterplot of pruned networks with export reactions (see main text) generated as described in **A**. Each point represents a different pruned network and the color of each point indicates the biomass reaction of that network. **C**. Same as **B** but colors indicate which set of nutrients the network was pruned with. **D**. Same as **B** but colors indicate growth rate of pruned network. **E-G**. Same as **B-D** but networks were pruned without export reactions (see main text). All pruned networks were derived from the universal string chemistry network with *A* = 2 and *L* = 5.

To assess the possibility that these results were just an artifact of the arbitrarily-chosen number of nutrients and biomass precursors, we investigated how the proportion of pruned reactions and connectivity of pruned networks change as the numbers of nutrients and biomass precursors vary (Figures S2 and S3). While the proportion of pruned reactions clearly decreases as the number of biomass precursors increases, as one might expect, it does not appear to be affected by the number of available nutrients. Figure S3 indicates that the metabolite to reaction ratio is always around 1 in pruned networks. These values are all slightly lower than those observed in real metabolic networks (see arrow in Figure 2), which likely reflects the fact that real metabolic networks must be capable of sustaining growth on multiple different environments, while the pruned networks are only required to sustain growth on one particular environment (see Methods). While we expect that the number of biomass precursors and nutrients may affect the structure of pruned networks in other more subtle ways, these findings support the idea that the results shown in Figure 4 do not depend on the number of biomass precursors or nutrients used during the pruning process.

## Discussion

We have created ARCHNET, a Python package capable of performing stoichiometric modeling on string chemistry networks of arbitrary size and complexity, and we devised an algorithm that identifies minimal metabolic networks necessary for converting a given set of environmental metabolites into a specific combination of “biomass” precursors within this framework. By running this pruning algorithm on many string chemistry networks, we found that the choice of the biomass metabolites wields much more influence over the structure of the minimal network than the choice of nutrients. Beyond this finding, our package could be used to further quantitatively explore any aspect of the complex relationship between metabolic network structure, environmental complexity, and biomass composition with minimal additional effort.

The biomass compositions of our string chemistry networks shed light on the processes underlying those in real metabolic networks. Many bacterial Genome-Scale Models (GEMs) are often created using “template” biomass reactions for key taxa, given the challenges of obtainingas opposed to having biomass compositions based on organism-specific experimental data [6,36]. While this is an open and fast-developing research area, one may wonder to what extent bacterial GEMs’ biomass reactions based on templates from a few organisms may affect the specificity of the model fluxes [36]. Our finding about the importance of biomass composition in string chemistry networks underscores the importance of careful reconstruction of biomass reactions in real metabolic networks [6,36,37].

The pruning algorithm is reminiscent of algorithms for identifying Elementary Flux Modes (EFMs) [38] and, identifying Minimal Balanced Pathways [17], and of certain gap-filling algorithms [39,40]. However, unlike EFMs, our pruned networks represent a single minimal set of reactions for transforming an arbitrary set of input metabolites into an arbitrary set of stoichiometrically constrained output metabolites, rather than a description of the entire steady state flux space. Gap-filling algorithms are increasingly developed and used to transform initial drafts of genome-scale metabolic reconstructions obtained from automated genome annotation into well-connected metabolic networks capable of producing the organism’s biomass from precursors, in a way that is compatible with experimental observations [40–44]. Gap-filling algorithms often approach this problem by adding to the initial network specific reactions from a large pool iteratively converging to an optimally gap-filled network [44–46]. Alternative algorithms have proposed carving the gap-filled network from a super-set of reactions [5]. Both gap-filling approaches bear some similarities to our pruning algorithm, suggesting that string chemistries could be used to simulate and further enhance these approaches, taking advantage of the tunable level of complexity of artificial networks, and on the complete knowledge of the underlying chemical universes.

Several previous studies used artificial chemistry as an avenue for addressing questions related to the origin of life or to general mathematical properties of biochemical networks [9–11,16,47]. Conversely, FBA has been applied mostly to the study of metabolic networks of real organisms [27–29]. There is likely great untapped potential available from combining the two approaches. In particular, the recent application of stoichiometric approaches to the study of early metabolism [49] and of ecosystem-level biochemical networks [50–52] could greatly benefit from additional creative usage of artificial chemistries. For example, the capacity to handle artificial string chemistries of arbitrary complexity using these same stoichiometric tools makes it possible to explore evolutionary processes and ecosystem-level metabolism under simulated scenarios in which the whole chemical universe is fully known. This will make it possible to shed light on the role of historical contingency and optimality principles in shaping the architecture of metabolic networks.

## Acknowledgements

This work was partially supported by the Directorates for Biological Sciences (BIO) and Geosciences (GEO) at the NSF and NASA under Agreements No. 80NSSC17K0295, 80NSSC17K0296 and 1724150 issued through the Astrobiology Program of the Science Mission Directorate. ARP was supported by a Howard Hughes Medical Institute Gilliam Fellowship and a National Academies of Sciences, Engineering, and Medicine Ford Foundation Predoctoral Fellowship. DM was supported by the National Institute of General Medical Sciences of the National Institutes of Health under award number T32GM100842.

---

**Figure S1.**
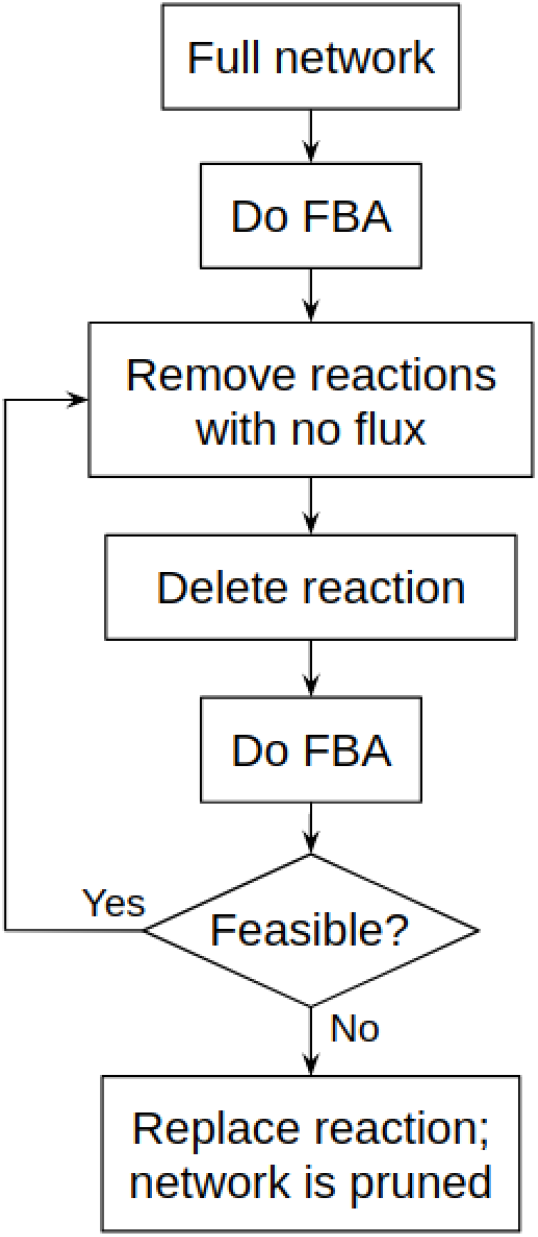
Description of the pruning algorithm.

**Figure S2.**
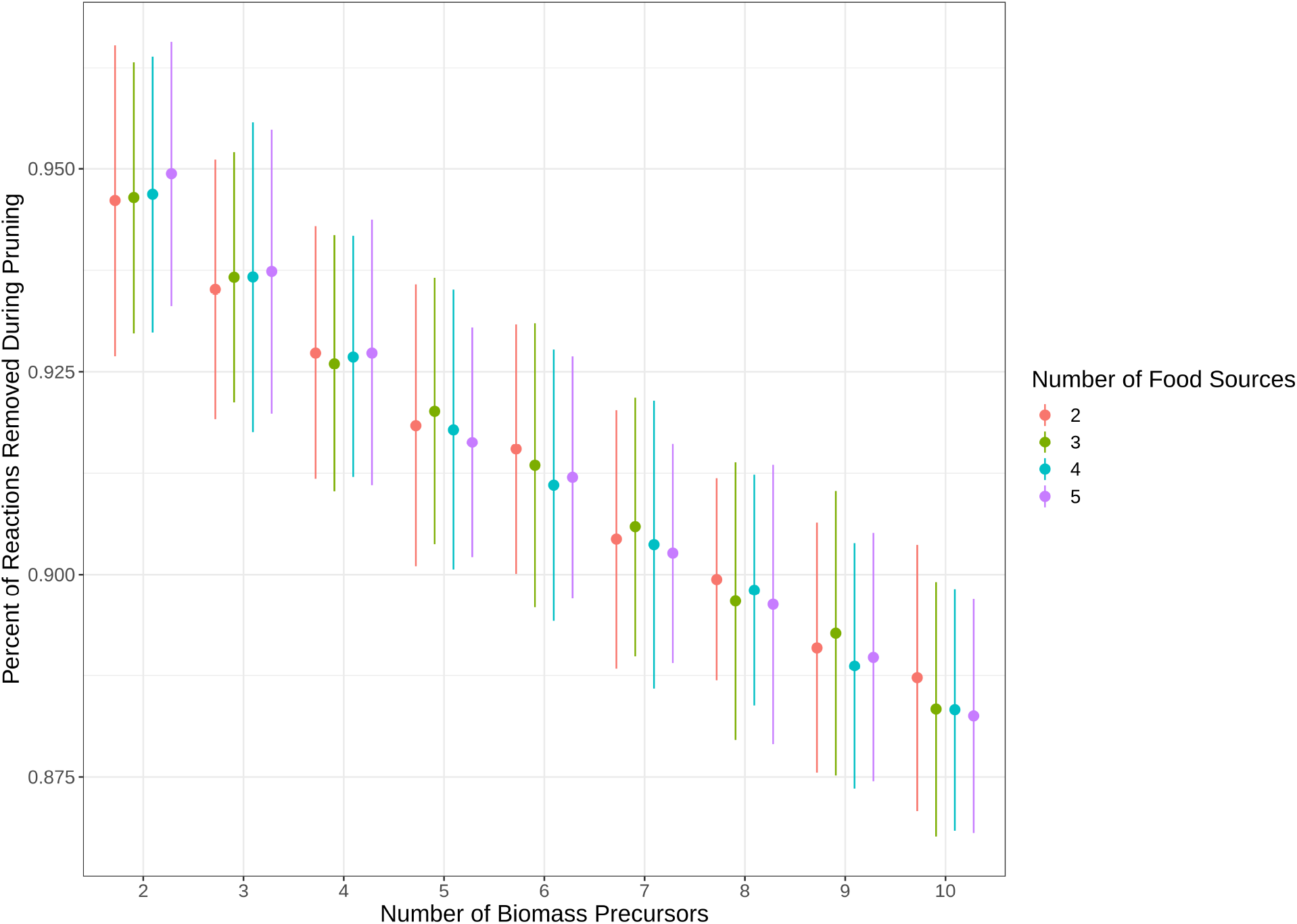
Number of biomass precursors affects the percentage of pruned reactions more than the number of nutrients. U2,5 was pruned with 100 different combinations of each number of food sources and biomass precursors shown on the graph. Each point is the mean pruned percentage with error bars indicating the standard deviation.

**Figure S3.**
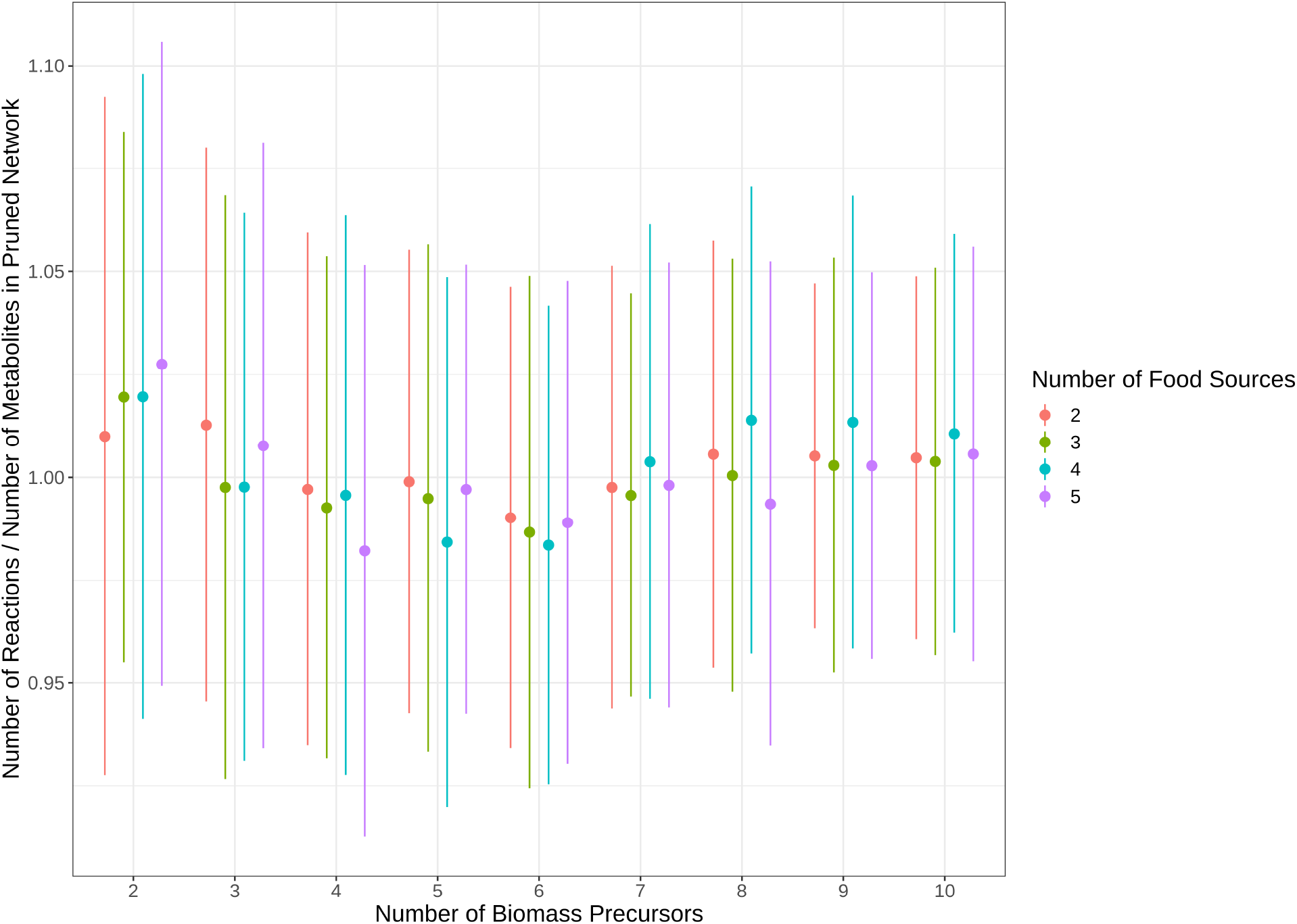
Number of biomass precursors affects the ratio of reactions to metabolites in pruned networks more than the number of nutrients. All networks were pruned from the chemical universe where *A* = 2 and *L* = 5 with export reactions allowed. Each point is the mean reaction-to-metabolite ratio with error bars indicating the standard deviation.

## References

1. Pál C, Papp B, Lercher MJ, Csermely P, Oliver SG, Hurst LD. Chance and necessity in the evolution of minimal metabolic networks. Nature. 2006;440: 667–670.

2. Barve A, Wagner A. A latent capacity for evolutionary innovation through exaptation in metabolic systems. Nature. 2013;500: 203–206.

3. Noor E, Eden E, Milo R, Alon U. Central carbon metabolism as a minimal biochemical walk between precursors for biomass and energy. Mol Cell. 2010;39: 809–820.

4. Ebenhöh O, Heinrich R. Evolutionary optimization of metabolic pathways. Theoretical reconstruction of the stoichiometry of ATP and NADH producing systems. Bull Math Biol. 2001. Available: https://link.springer.com/article/10.1006/bulm.2000.0197

5. Machado D, Andrejev S, Tramontano M, Patil KR. Fast automated reconstruction of genome-scale metabolic models for microbial species and communities. Nucleic Acids Res. 2018;46: 7542–7553.

6. Henry CS, DeJongh M, Best AA, Frybarger PM, Linsay B, Stevens RL. High-throughput generation, optimization and analysis of genome-scale metabolic models. Nat Biotechnol. 2010;28: 977–982.

7. Raymond J, Segrè D. The effect of oxygen on biochemical networks and the evolution of complex life. Science. 2006;311: 1764–1767.

8. Handorf T, Ebenhöh O, Heinrich R. Expanding metabolic networks: scopes of compounds, robustness, and evolution. J Mol Evol. 2005;61: 498–512.

9. Banzhaf W, Yamamoto L. Artificial Chemistries. MIT Press; 2015.

10. Benkö G, Flamm C, Stadler PF. A graph-based toy model of chemistry. J Chem Inf Comput Sci. 2003;43: 1085–1093.

11. Kauffman SA, Member of the Santa Fe Institute and Professor of Biochemistry Stuart A Kauffman. The Origins of Order: Self-organization and Selection in Evolution. Oxford University Press; 1993.

12. Guseva E, Zuckermann RN, Dill KA. Foldamer hypothesis for the growth and sequence differentiation of prebiotic polymers. Proc Natl Acad Sci U S A. 2017;114: E7460–E7468.

13. Friedlander T, Mayo AE, Tlusty T, Alon U. Evolution of bow-tie architectures in biology. PLoS Comput Biol. 2015;11: e1004055.

14. Fontana W, Buss LW. What would be conserved if “the tape were played twice”? Proc Natl Acad Sci U S A. 1994;91: 757–761.

15. Pfeiffer T, Soyer OS, Bonhoeffer S. The evolution of connectivity in metabolic networks. PLoS Biol. 2005;3: e228.

16. Fontana W, Buss LW. “The arrival of the fittest”: Toward a theory of biological organization. Bull Math Biol. 1994;56: 1–64.

17. Riehl WJ, Krapivsky PL, Redner S, Segrè D. Signatures of arithmetic simplicity in metabolic network architecture. PLoS Comput Biol. 2010;6: e1000725.

18. Soyer OS, Pfeiffer T. Evolution under fluctuating environments explains observed robustness in metabolic networks. PLoS Comput Biol. 2010;6. doi:10.1371/journal.pcbi.1000907

19. Weininger D. SMILES, a chemical language and information system. 1. Introduction to methodology and encoding rules. J Chem Inf Comput Sci. 1988;28: 31–36.

20. Arús-Pous J, Johansson SV, Prykhodko O, Bjerrum EJ, Tyrchan C, Reymond J-L, et al. Randomized SMILES strings improve the quality of molecular generative models. J Cheminform. 2019;11: 71.

21. Lin T-S, Coley CW, Mochigase H, Beech HK, Wang W, Wang Z, et al. BigSMILES: A Structurally-Based Line Notation for Describing Macromolecules. ACS Cent Sci. 2019;5: 1523–1531.

22. Lee AA, Yang Q, Sresht V, Bolgar P, Hou X, Klug-McLeod JL, et al. Molecular Transformer unifies reaction prediction and retrosynthesis across pharma chemical space. Chem Commun. 2019;55: 12152–12155.

23. Heirendt L, Arreckx S, Pfau T, Mendoza SN, Richelle A, Heinken A, et al. Creation and analysis of biochemical constraint-based models using the COBRA Toolbox v.3.0. Nat Protoc. 2019;14: 639–702.

24. Ebrahim A, Lerman JA, Palsson BO, Hyduke DR. COBRApy: COnstraints-Based Reconstruction and Analysis for Python. BMC Syst Biol. 2013;7: 74.

25. Gottstein W, Olivier BG, Bruggeman FJ, Teusink B. Constraint-based stoichiometric modelling from single organisms to microbial communities. J R Soc Interface. 2016;13. doi:10.1098/rsif.2016.0627

26. O’Brien EJ, Monk JM, Palsson BO. Using Genome-scale Models to Predict Biological Capabilities. Cell. 2015;161: 971–987.

27. Orth JD, Thiele I, Palsson BØ. What is flux balance analysis? Nat Biotechnol. 2010;28: 245–248.

28. Gu C, Kim GB, Kim WJ, Kim HU, Lee SY. Current status and applications of genome-scale metabolic models. Genome Biol. 2019;20: 121.

29. Kauffman KJ, Prakash P, Edwards JS. Advances in flux balance analysis. Curr Opin Biotechnol. 2003;14: 491–496.

30. Chang R. Physical Chemistry for the Biosciences. University Science Books; 2005.

31. Laidler KJ, Glasstone S. Rate, order and molecularity in chemical kinetics. J Chem Educ. 1948;25: 383.

32. Compton RG, Bamford CH, Tipper† CFH. The Theory of Kinetics. Elsevier; 2012.

33. Hucka M, Finney A, Sauro HM, Bolouri H, Doyle JC, Kitano H, et al. The systems biology markup language (SBML): a medium for representation and exchange of biochemical network models. Bioinformatics. 2003;19: 524–531.

34. Pacheco AR, Moel M, Segrè D. Costless metabolic secretions as drivers of interspecies interactions in microbial ecosystems. Nat Commun. 2019;10: 103.

35. McInnes L, Healy J, Melville J. UMAP: Uniform Manifold Approximation and Projection for Dimension Reduction. arXiv [stat.ML]. 2018. Available: http://arxiv.org/abs/1802.03426

36. Xavier JC, Patil KR, Rocha I. Integration of Biomass Formulations of Genome-Scale Metabolic Models with Experimental Data Reveals Universally Essential Cofactors in Prokaryotes. Metab Eng. 2017;39: 200–208.

37. Lachance J-C, Lloyd CJ, Monk JM, Yang L, Sastry AV, Seif Y, et al. BOFdat: Generating biomass objective functions for genome-scale metabolic models from experimental data. PLoS Comput Biol. 2019;15: e1006971.

38. Schuster S, Fell DA, Dandekar T. A general definition of metabolic pathways useful for systematic organization and analysis of complex metabolic networks. Nat Biotechnol. 2000;18: 326–332.

39. Orth JD, Palsson BØ. Systematizing the generation of missing metabolic knowledge. Biotechnol Bioeng. 2010;107: 403–412.

40. Satish Kumar V, Dasika MS, Maranas CD. Optimization based automated curation of metabolic reconstructions. BMC Bioinformatics. 2007;8: 212.

41. Thiele I, Vlassis N, Fleming RMT. fastGapFill: efficient gap filling in metabolic networks. Bioinformatics. 2014;30: 2529–2531.

42. Prigent S, Frioux C, Dittami SM, Thiele S, Larhlimi A, Collet G, et al. Meneco, a Topology-Based Gap-Filling Tool Applicable to Degraded Genome-Wide Metabolic Networks. PLoS Comput Biol. 2017;13: e1005276.

43. Christian N, May P, Kempa S, Handorf T, Ebenhöh O. An integrative approach towards completing genome-scale metabolic networks. Mol Biosyst. 2009;5: 1889–1903.

44. Vitkin E, Shlomi T. MIRAGE: a functional genomics-based approach for metabolic network model reconstruction and its application to cyanobacteria networks. Genome Biol. 2012;13: R111.

45. Reed JL, Patel TR, Chen KH, Joyce AR, Applebee MK, Herring CD, et al. Systems approach to refining genome annotation. Proc Natl Acad Sci U S A. 2006;103: 17480–17484.

46. Pharkya P, Burgard AP, Maranas CD. OptStrain: a computational framework for redesign of microbial production systems. Genome Res. 2004;14: 2367–2376.

47. Peng Z, Plum AM, Gagrani P, Baum DA. An ecological framework for the analysis of prebiotic chemical reaction networks. J Theor Biol. 2020; 110451.

48. Goldford JE, Hartman H, Marsland R 3rd, Segrè D. Environmental boundary conditions for the origin of life converge to an organo-sulfur metabolism. Nat Ecol Evol. 2019. doi:10.1038/s41559-019-1018-8

49. Goldford JE, Segrè D. Modern views of ancient metabolic networks. Current Opinion in Systems Biology. 2018. Available: https://www.sciencedirect.com/science/article/pii/S2452310017302196

50. Carlson RP, Beck AE, Phalak P, Fields MW, Gedeon T, Hanley L, et al. Competitive resource allocation to metabolic pathways contributes to overflow metabolisms and emergent properties in cross-feeding microbial consortia. Biochem Soc Trans. 2018;46: 269–284.

51. Klitgord N, Segrè D. Environments that induce synthetic microbial ecosystems. PLoS Comput Biol. 2010;6: e1001002.

52. Harcombe WR, Riehl WJ, Dukovski I, Granger BR, Betts A, Lang AH, et al. Metabolic resource allocation in individual microbes determines ecosystem interactions and spatial dynamics. Cell Rep. 2014;7: 1104–1115.

